# No change in oxytocin level before a human intergroup competition: study among Polish players before rugby and handball matches

**DOI:** 10.1101/2020.11.30.403592

**Authors:** Slawomir Koziel, Marek Kociuba, Zofia Ignasiak, Andrzej Rokita, Ireneusz Cichy, Andrzej Dutkowski, Marcin Ściślak, Katarzyna Kochan, Anna Sebastjan, Anna Spinek, Daria Lorek, Raja Chakraborty, Barry Bogin

## Abstract

**Objectives:** The aim of the present study was to assess the changes in urinary oxytocin concentration during the period between five days before, and on the day of match, among rugby and handball players.

**Methods:** The study used a repeated measures design with the relative oxytocin level as the outcome variable measured at two subsequent points of time, viz., on five days before as well as on the days of matches. Nine male rugby players with a mean age of 27.62 years (SD = 4.21) and 18 male handball players with a mean age of 17.03 years (SD = 0.57) participated. Urinary oxytocin level was measured by ELISA immunoassay as a ratio to the concentration of creatinine [mg/ml] measured through colorimetric detection. Differences in oxytocin level were assessed by ANOVA with repeated measurements.

**Results:** The OT/CRE levels significantly differed between the type of player (rugby or handball) but not between times of measurements. Tukey’s post-hoc tests revealed that significant differences were only between OT/CRE level in a day of match in rugby players and in 5 days before match in handball players (p<0.05).

**Conclusion:** There was no change in oxytocin levels during the time periods between five days before and on the day of a match, in either of the two kinds of players. The change in oxytocin might be traceable during the match but not before a match and thus perhaps depends on a more subtle context of competition, but not on the assumption of competition.

## Introduction

Oxytocin (OT), the endogenous neuropeptide performing peripheral role as a hormone, is mostly known for facilitating uterine contractions during labour, facilitating maternal behaviour [1], and also for initiating parenting behaviour [2]. In addition, oxytocin has a broader role exerting modifying effects on human social cognition and behaviour [3] that increase benefits of social interaction and promote social approach and affiliation [4], build trust [5], attachment [6] and group cohesion [7,8] in humans and non-human animals [1.9-11]. The critical and intricate role of oxytocin has been implicated in the formation and maintenance of social groups by modifying several behaviours and human cooperative traits that are important in effectively functioning in a group [12-14]. However, there are recent reports that administration of oxytocin increased aggressive behaviour [15] and that oxytocin receptor gene polymorphism has association with aggression [16] that might be important in combat with out-group enemies. According to a theoretical framework, oxytocin modulates social behavior by means of increasing the salience of social stimuli and perhaps promotes a wide range of emotions and behaviors and not merely the positive and affiliative ones [17]. Therefore, if indeed oxytocin increases the salience of social agents, a plausible assumption is that it will increase aggressive reactions in competitive situations involving aggressive provocations [15]. In a study of intergroup conflict in wild chimpanzees Samuni et al. [18] reported that oxytocin levels were elevated immediately before and during intergroup conflict compared with controls.

Human inter-group conflicts are part of our evolutionary history and success in such conflicts depends upon several adaptive mechanisms. Sports and games sometimes posit on humans a challenge situation that mimics inter-group conflict, as well as intra-group coordination to win over the opponent [19]. Sporting activities are very often chosen as a proxy to human combative situations. The sport competition model may involve several adaptive mechanisms, including aggression, and this model has been in studies related to endocrine response or prenatal hormone exposures [20,21]. On the other hand, there is evidence for prosocial behavior in enhancing team performance in sport [22]. In a review, after considering several previous evidences, it was proposed that oxytocin could be the bridging link between certain kinds of empathy as well as emotion transfer and enhanced group performance in team sports (for detailed review see: Pepping and Timmermans 2012) [23].

Success in sports has been often linked with motivation and positive emotions [24]. Prosocial behaviours during a game, such as the high-five, the fist-pump, and the group hug, remain staple elements of success in sporting life [25]. These behaviours may enhance oxytocin [26]. But it is not known if such increase also occurs as a preparatory mechanism for a motivational adaptation among elite sports persons before a match. Pre-match elevated oxytocin might be important for soothing of stress and enhancing empathy and greater receptivity for collective emotions for teammates or rivalry against opponents.

The aim of the present study was to analyse the changes in urinary oxytocin concentration between 5 days before, and on the day of match, in rugby and handball players in Wroclaw, Poland. It was hypothesized that oxytocin would rise before the match as an adaptive preparation for a challenging situation and building capacity for enhanced team cooperation. The present study chose rugby and handball matches as these are known to regularly generate high level of competitive excitement both among the players and the spectators. Rugby involves high level of contact, aggressive interactions, and hence high prevalence of injuries; handball on the other hand requires specialized team coordination [27]. In this study, these two kinds of sports were chosen to proxy high aggressive competition and sociability, respectively.

## Materials and Methods

### Study design, participants, and settings

This study followed a repeated measures design of the outcome variable (oxytocin) at two subsequent points of time. Nine male rugby players with a mean age of 27.62 years (SD = 4.21), participating in games of regional rugby league (Lower Silesia), and 18 male handball players with a mean age of 17.03 years (SD = 0.57), taking part in games of academic league, were included in the study. All participants provided informed consent before participation. The samples for study (urine) were collected by the participants on their own at their respective homes. They collected the morning samples of urine 5 days before and on the day of match. For this purpose, sterile plastic containers (100 ml) were distributed to each participant before the day of collecting sample. They were asked to collect the samples of the first urine in one sterile container after waking up in the morning. After collection of these containers from the participants on the same day, each sample was stored in two sterile plastic tubes (30 ml) at minus 20° Celsius temperature till the laboratory analyses were conducted. The rugby players were followed for 3 matches (during April and May, 2017), whereas handball players for 2 matches (during October, 2017). Together, the 112 urine samples were collected, 51 from rugby and 61 from handball players.

### Sample analyses

After thawing, the samples were centrifuged at 1500 × g for 10 minutes at 4° Celsius temperature and supernatants were collected. Oxytocin concentration was estimated by ELISA immunoassay (Catalog #: ADI-901-153A; Enzo Life Science, USA). Creatinine (Cr) concentration was measured with the Creatinine Colorimetric Detection Kit (Catalog #: ADI-907-030A; Enzo Life Science, USA). Both analyses were conducted according to the assay procedures described in provided manuals. All standards and samples in both assays were run in duplicate. Oxytocin concentration in urine samples was expressed as a ratio of oxytocin to creatinine (oxytocin pg/mg creatinine) and used in further calculations. Analyses were conducted in a laboratory of Institute of Immunology and Experimental Therapy, Polish Academy of Sciences by two trained members of the institute (AS, DL).

### Statistics

Statistics of mean, standard deviation (SD), median, maximum-, and minimum values were used to describe the measures of anthropometry as well as oxytocin-to-creatinine level. Differences in ratios of oxytocin, between 5 days before and a day of match, were assessed by ANOVA with repeated measurements, where appropriate team and repeated estimation of oxytocin were factors. *Post hoc* comparison was done of the means of Tukey’s Test for unequal sample size. All calculations were done using STATISTICA 13.1 [28].

### Ethical approval

The study protocol has been approved by the Senate Ethical Committee Scientific Research of the University School of Physical Education in Wroclaw. Although the study used non-invasive measures, the ethical guidelines as laid down by the Helsinki Declaration was sincerely adhered to [29].

## Results

**Table 1** shows the mean age, height, weight and BMI in Rugby and Handball players and the difference in these measures between these two groups of players as assessed by student t test.

**Table 1.** Descriptive statistics of age and body dimensions of the players

|  |  | Age |  | Height |  | Weight |  | BMI |  |
| --- | --- | --- | --- | --- | --- | --- | --- | --- | --- |
|  | N | Mean | SD | Mean | SD | Mean | SD | Mean | SD |
| Rugby | 9 | 27.6 | 4.2 | 181.3 | 6.4 | 90.0 | 7.6 | 27.3 | 1.2 |
| Handball | 18 | 17.8 | 0.60 | 183.3 | 6.1 | 83.4 | 12.8 | 24.7 | 3.0 |
| t-test |  | p<0.001 |  | p>0.05 |  | p>0.05 |  | p<0.05 |  |

There were significant differences in age (p<0.001) and BMI (p<0.05), but not in height and weight. The rugby players were older and had higher BMI than the handball players.

**Table 2** presents the descriptive statistics of the ratio of oxytocin-to-creatinine levels [OT/CRE, pg/mg] for each subject. For each rugby player six estimations were done, whereas for handball players only four.

**Table 2.** Descriptive statistics of the ratio of oxytocin-to-creatinine [pg/mg] for individual players.

| Players' number | N-urine samples | Mean | Median | minimum | maximum | SD |
| --- | --- | --- | --- | --- | --- | --- |
| Rugby (N = 52) |  |  |  |  |  |  |
| 2 | 6 | 437.6 | 383.6 | 686.2 | 238.4 | 188.9 |
| 3 | 6 | 608.6 | 549.4 | 879.6 | 465.8 | 162.6 |
| 5 | 6 | 684.9 | 626.4 | 943.1 | 511.0 | 176.0 |
| 7 | 6 | 648.0 | 534.8 | 1073.0 | 383.9 | 262.5 |
| 8 | 6 | 665.7 | 639.4 | 843.4 | 535.0 | 108.0 |
| 9 | 6 | 663.6 | 664.1 | 738.3 | 575.2 | 58.7 |
| 10 | 6 | 864.0 | 848.8 | 1054.7 | 721.7 | 143.6 |
| 11 | 6 | 724.8 | 714.7 | 941.2 | 477.8 | 172.7 |
| 12 | 4 | 514.0 | 547.8 | 562.9 | 397.5 | 78.1 |
| All | 52 | 650.7 | 635.0 | 1073.0 | 238.4 | 189.4 |
| Handball (N = 61) |  |  |  |  |  |  |
| 1 | 2 | 902.2 | 902.2 | 891.7 | 912.8 | 14.9 |
| 2 | 3 | 760.7 | 773.1 | 728.3 | 780.7 | 28.3 |
| 3 | 3 | 629.5 | 651.5 | 494.0 | 743.0 | 125.9 |
| 4 | 3 | 839.5 | 784.9 | 782.4 | 951.1 | 96.7 |
| 6 | 4 | 597.8 | 611.9 | 464.0 | 703.5 | 106.2 |
| 7 | 4 | 703.7 | 710.3 | 630.0 | 764.3 | 62.1 |
| 8 | 4 | 614.9 | 645.9 | 454.1 | 713.7 | 112.3 |
| 9 | 4 | 777.9 | 812.9 | 549.8 | 935.8 | 171.3 |
| 10 | 4 | 565.7 | 575.9 | 401.5 | 709.3 | 130.9 |
| 11 | 3 | 637.0 | 678.2 | 500.3 | 732.5 | 121.4 |
| 12 | 4 | 737.0 | 778.7 | 575.5 | 815.0 | 113.0 |
| 13 | 2 | 687.3 | 687.3 | 685.0 | 689.5 | 3.2 |
| 14 | 2 | 918.4 | 918.4 | 726.8 | 1110.1 | 271.0 |
| 15 | 4 | 857.6 | 754.0 | 607.0 | 1315.5 | 313.1 |
| 16 | 4 | 941.5 | 935.3 | 745.4 | 1149.9 | 204.6 |
| 17 | 4 | 891.0 | 880.7 | 757.7 | 1044.8 | 120.3 |
| 18 | 3 | 773.4 | 815.5 | 666.3 | 838.5 | 93.5 |
| 21 | 4 | 747.5 | 697.1 | 619.2 | 976.4 | 167.5 |
| All players | 61 | 748.8 | 742.6 | 401.5 | 1315.5 | 172.5 |

Results of ANOVA revealed that only the type of player (rugby or handball) significantly differentiated OT/CRE levels, but no effect of repeated measurements (**Table 3**). Tukey’s post-hoc tests showed only differences between OT/CRE level in a day of match in rugby players and OT/CRE level in 5 days before match in handball players (p<0.05).

**Table 3.** Results of ANOVA with repeated measurements of the oxytocin-to-creatinine ratio [pg OT/mg Cr] in 5 days and in a day of match among rugby and handball players

|  | F | p |
| --- | --- | --- |
| Players | 6.63 | <0.05 |
| Repeated measurements | 1.17 | n.s. |
| Interaction | 0.26 | n.s. |

**Figure 1** demonstrates the changes in the OT/CRE levels in each player between five days before match and on the day of match. In this graph no definite pattern or regularity was observed indicating any difference in ratios 5 days before and on a day of match.

**Figure 1.**
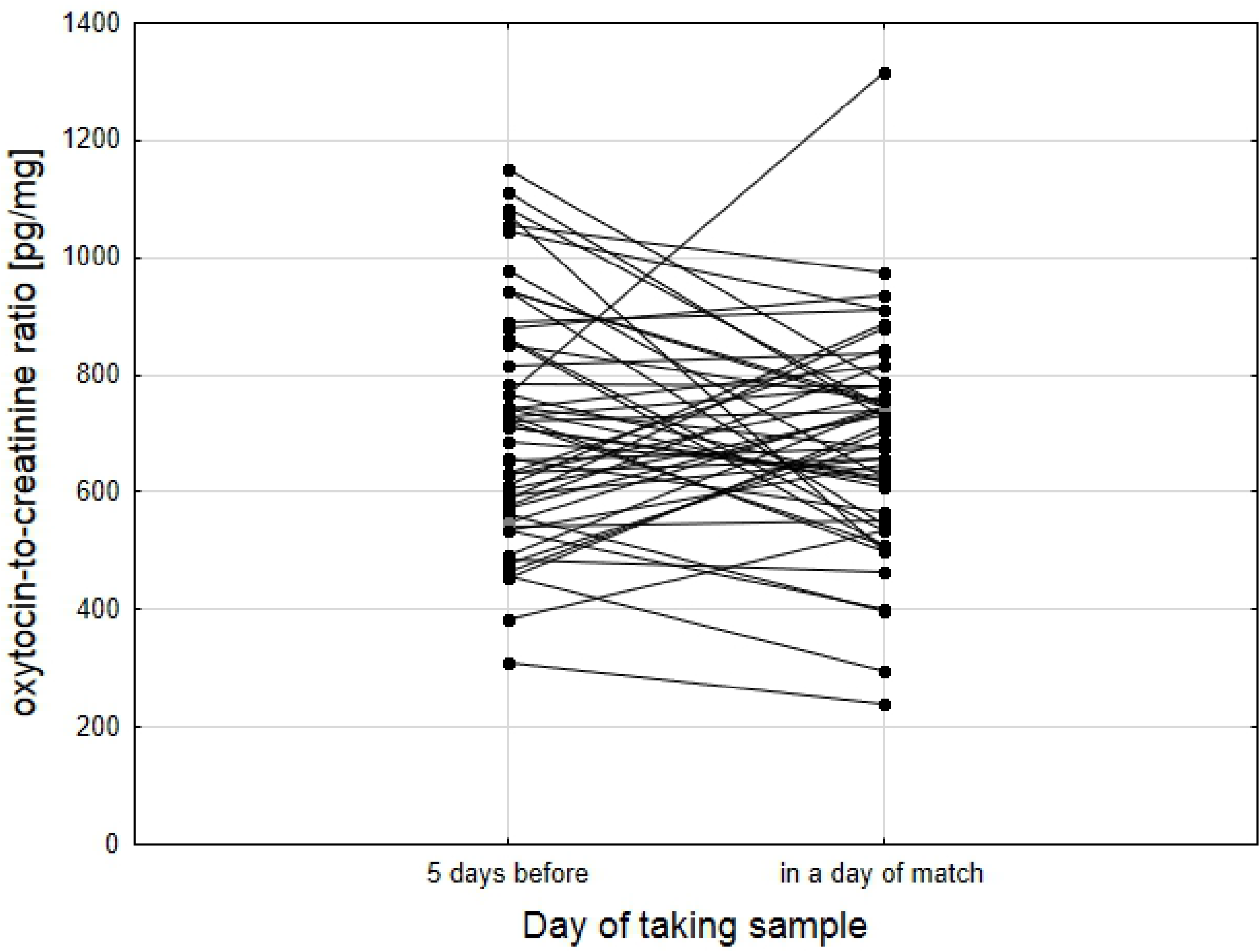
Changes of individual level of oxytocin-to-creatinine ratio [pg/mg] in all players estimated 5 days before and in a day of the match.

## Discussion

The effects of oxytocin on adaptive social behaviour have been the topic of increasing scientific interest in recent years, both in animal and human studies. In the present study, we investigated whether urinary oxytocin level among elite players would rise up on the day, as compared to five days, before highly competitive football and handball matches.

Most research evidence was obtained in laboratory settings using intergroup social-dilemma games and focused on human male participants. Only a handful of studies have studied intergroup contexts in captive or wild non-human animals. In a study among wild chimpanzees, during intergroup conflicts in natural habitat, a significantly higher urinary oxytocin level was recorded in both sexes immediately before and during the intergroup conflicts (compared to controls). Both anticipatory response to, and participation in the intergroup conflicts involved high urinary oxytocin levels compared to control conditions [18, 30].

During intergroup conflict, elevated levels of both cortisol and oxytocin are expected to have adaptive importance. Whereas the former facilitates rapid production of energy required for combat or flight [31], the latter may promote the essential in-group cohesion leading to cooperative responses [7,18]. Studies on both humans and chimpanzees indicate an oxytocinergic system involvement during intergroup conflict [7,18]. It is widely believed that sports and games could closely imitate a real-life inter-group conflict and intra-group cohesion among humans [19]. It was therefore hypothesised that similar effect could also be visible shortly before participation in highly competitive games. In Poland, rugby is played by a very few teams and the handball players selected were relatively top ranking. Therefore, it could be assumed that both types of players participating in this study were dedicated to their games. Thus, the players were expected to develop their ability to perform at a high level by several years of practice and adjustment to the specific requirements of the game. They were, therefore, expected to show adaptive physiological changes, such as raised oxytocin, that might support them in performance, shortly before a match. However, the results of the present study did not provide such evidence showing no definite pattern or regularity of changes in oxytocin level from five days before and on a day of match. In both types of games there were no significant differences between those two time points. The result was identical even when data for all players in two games were analysed together.

There are several probable reasons for the ‘negative’ result. With growing number of investigations on the effects of oxytocin on behaviour, cognition, and neuropsychiatry, it is becoming clear that its functions are far more complex than initially was presumed [9,10]. For example, a more recent study among chimpanzees did not find any association between oxytocin level and being in proximity to the territorial boundary, although it is known that being near the territorial boundary between social groups increases the chances of intergroup conflict [32]. The researchers suggested that exposure to a stressor, such as and intergroup fight, might not be the main impulse for oxytocin secretion in chimpanzees. Oxytocin release during intergroup conflict as observed by Samuni et al. [18] was thus postulated to be triggered by the social context that was linked with the stressor rather than the stressor itself [32]. Accumulated evidence from several studies indicates that social contexts and individual factors, such as sex, early experience, or health, influence oxytocinergic effects [1,9]. Perhaps, the context before actual commencement of the matches of our participant players influenced their oxytocin production. Just being near the ‘boundary’ for a match, that is, near the day of a match, is not the trigger. Rather, physical-aggressive contact during the match would raise oxytocin levels. Unfortunately, we did not measure OT/CRE ratios during or after the game.

Several studies, e.g., Striepens et al., [33], suggested that oxytocin mediates behaviors which are mainly prosocial. Nonetheless, other studies from animal research contradicted this by showing that oxytocin can also enhance anxiety [34]. Similarly, among humans, recent studies demonstrated its effects that might promote conflict rather than cohesion. For example, oxytocin was found to increase envy and gloating [35], decrease trust and the inclination to cooperate in individuals with borderline personality disorder [9], and facilitate out-group derogation [8]. Thus, although oxytocin is widely viewed as a prosocial compound, it may also promote antisocial responses, thus suggesting a context-dependent effect [36]. Thus, it seemed that oxytocin level is not raised, at least, at this stage (before a few hours of match), but maybe just before game or during game. Instead, prosocial behaviours, such as mutual touching and hugging among teammates during the game (context) might have increased the oxytocin level [25,26]. Again, we did not collect samples during or after the match.

Although sports and games could suitable proxies for real life competitions [21], there are certain differences between sport, game, and real life. It would be ideal to measure oxytocin in soldiers at war, as was done for fighting chimpanzees. The present study, however, showed that although some oxytocinergic response during preparatory period of a competitive sport was expected, it might not have been activated even a few hours before the match. Perhaps, the social bonds between the members of a group of chimpanzees are much stronger than among teammates in a game. The chimpanzees were brought up together in their social group and usually do not have opportunities for social cohesion with strangers. In contrast, humans very often do have these opportunities. Therefore, social networks and requirements for cooperation and affiliation are quite different between these two species. These differences may result in markedly divergent patterns of oxytocin release.

The social context immediately before a conflict to protect territory among the chimpanzee is expected to be very different from that of a human sports person before playing in a match. The game was only a part of their life but for chimpanzees the fights are to protect their entire world. Studies on animal models have also demonstrated that social context might modify the regulating effect of oxytocin on social play [37]. In conclusion, the present study showed that the oxytocin level in elite sportspersons did not increase with the nearing of a match. Perhaps the increase might be noted in a more critical context, such as, during the game.

## Acknowledgements

The authors are especially grateful to all rugby and handball players for their participation in the study.

## Author contributions

SK - designed the study, made the analysis and prepared the draft and checked the final version; MK - collected the urine samples, prepared samples for analysis, build the database, ZI, AR, IC, AD, MŚ, KK, AS - recruited and instructed participants, conducted measurements and prepare database, prepare first draft; AS, DL - carried out biochemical analysis, prepared part of the draft, checked final draft; RC, BB - prepared and edited the final draft, collected literature, corrected statistical analysis.

## References

1. Rilling JK, DeMarco AC, Hackett PD, et al. Effects of intranasal oxytocin and vasopressin on cooperativebehavior and associated brain activity in men. Psychoneuroendocrinol 2012; 37(4):447–461. https://doi.org/10.1016/j.psyneuen.2011.07.013

2. Gordon I, Zagoory-Sharon O, Leckman J, et al. Oxytocin and the development of parenting in humans. Biol psychiatr 2010; 68:377–382. https://doi.org/10.1016/j.biopsych.2010.02.005

3. Bartz JA, Zaki J, Bolger N, et al. Social effects of oxytocin in humans: context and person matter. Trends Cogn Sci 2011; 15:301–309. https://doi.org/10.1016/j.tics.2011.05.002

4. Heinrichs M, Domes G. Neuropeptides and social behaviour: effects of and vasopressin in humans. Prog Brain Res 2008; 170:337–350. https://doi.org/10.1016/S0079-6123(08)00428-7

5. Kosfeld M, Heinrichs M, Zak PJ, et al. Oxytocin increases trust in humans. Nature 2005; 435:673–676. https://doi.org/10.1038/nature03701

6. Donaldson ZR, Young LJ. Oxytocin, vasopressin, and the neurogenetics of sociality. Science 2008; 322:900–4. https://doi.org/10.1126/science.1158668

7. De Dreu CKW, Greer LL, Handgraaf MJJ, et al. The neuropeptide oxytocin regulates parochial altruism in intergroup conflict among humans. Science 2010; 328:1408–1411. https://doi.org/10.1126/science.1189047

8. De Dreu CKW, Greer LL, Van Kleef GA, et al. Oxytocin promotes human ethnocentrism. Proc Natl Acad Sci 2011; 108:1262–1266. https://doi.org/10.1073/pnas.1015316108

9. Bartz J, Simeon D, Hamilton H, et al. Oxytocin can hinder trust and cooperation in borderline personality disorder. Soc Cogn Affect Neurosci 2011; 6:556–563.

10. Chang SWC, Brent LJN, Adams, GK, et al. Neuroethology of primate social behaviour. Proc Natl Acad Sci 2013; 110:10387–10394. https://doi.org/10.1073/pnas.1301213110

11. Neumann ID, Slattery DA. Oxytocin in general anxiety and social fear: a translational approach. Biol Psychiatry Oxytocin and Psychiatry: From DNA to Social Behavior 2016; 79:213–221. https://doi.org/10.1016/j.biopsych.2015.06.004

12. De Dreu CKW. Oxytocin modulates cooperation within and competition between groups: An integrative review and research agenda. Horm Behav 2012; 61:419–428. https://doi.org/10.1016/i.yhbeh.2011.12.009

13. Choi J-K, Bowles S. The coevolution of parochial altruism and war. Science 2007; 318:636–640. https://doi.org/10.1126/science.1144237

14. Bowles S. Did warfare among ancestral hunter-gatherers affect the evolution of human social behaviors? Science 2009; 324:1293–1298. https://doi.org/10.1126/science.1168112

15. Ne’eman R, Perach-Barzilay N, Fischer-Shofty M, et al. Intranasal administration of increases human aggressive behaviour. Horm Behav 2016; 80:125–130. https;//doi.org/10.1016/j.yhbeh.2016.01.015

16. Butovskaya M, Rostovtseva V, Butovskaya P. et al. Oxytocin receptor gene polymorphism (rs53576) and digit ratio associates with aggression: comparison in seven ethnic groups. J Physiol Anthropol 2020; 39. https://doi.org/10.1186/s40101-020-00232-y

17. Shamay-Tsoory SG, Abu-Akel A. The social salience hypothesis of oxytocin. Biol Psychiatry 2016; 66: 864–870. https://doi.org/10.1016/j.biopsych.2015.07.020

18. Samuni L, Preis A, Mundry R, et al. Oxytocin reactivity during intergroup conflict in wild chimpanzees. Proc Natl Acad Sci 2016; 114:268–273 https://doi.org/10.1073/pnas.1616812114

19. Muñoz-Reyes JA, Polo P, Valenzuela N, et al. The male warrior hypothesis: testosterone-related cooperation and aggression in the context of intergroup conflict. Sci Rep 2020; 10. https://doi.org/10.1038/s41598-019-57259-0

20. Bogin B, Hermanussen M, Blum WF, Aßmann C. Sex, Sport, IGF-1 and the community effect in height hypothesis. Int J Environ Res Pub Health 2016: 12(5), 4816–4832. http://www.mdpi.com/1660-4601/12/5/4816

21. Longman D, Stock JT, Wells JCK. Digit ratio (2D:4D) and rowing ergometer performance in males and females, Am J Phys Anthropol 2011; 44:337–341. https://doi.org/1002/ajpa.21407

22. Moll T, Jordet G, Pepping GJ. Emotional contagion in soccer penalty shootouts: celebration of individual success is associated with ultimate team success. J Sports Sci 2010; 28:983–992. https://doi.org/10.1080/02640414.2010.484068

23. Pepping G, Timmermans EJ. Oxytocin and the Biopsychology of Performance in Team Sports. Sci World J 2012; https://doi.org/10.1100/2012/567363

24. McCarthy PJ. Positive emotion in sport performance: current status and future directions. Int Rev Sport Exer Psychol 2011; 4:50–69. https://doi.org/10.1080/1750984X.2011.560955

25. Kraus MW, Huang C, Keltner D. Tactile communication, cooperation, and performance: an ethological study of the NBA. Emotion 2010; 10:745–749. https://doi.org10.1037/a0019382

26. Uvnäs-Moberg K, Handlin L, Petersson M. Self-soothing behaviors with particular reference to oxytocin release induced by non-noxious sensory stimulation. Front Psychol 2015; 5:1529. https://doi.org/10.3389/fpsyg.2014.01529

27. Cummins C, Orr R. Analysis of physical collisions in elite national rugby league match play. Int J Sports Physiol Perform. 2015; 10:732–9. https://doi.org/10.1123/iispp.2014-0541

28. TIBCO Software Inc. (2017). Statistica (data analysis software system), version 13. http://statistica.io.

29. Goodyear MDE, Krleza-Jeric K, Lemmens T. The Declaration of Helsinki. Br Med J 2007; 335:624–625. doi: 10.1136/bmj.39339.610000.BE

30. Samuni L, Preis A, Deschner T, et al. Reward of labor coordination and hunting success in wild chimpanzees. Commun Biol 2018; 1:138. https://doi.org/10.1038/s42003-018-0142-3

31. McEwen BS. Physiology and neurobiology of stress and adaptation: central role of the brain. Physiol Rev 2007; 87: 873–904. https://doi.org/10.1152/physrev.00041.2006

32. Samuni L, Preis A, Deschner T et al. Cortisol and show independent activity during chimpanzee intergroup conflict. Psychoneuroendocrinol 2019; 104: 165–173 https://doi.org/10.1016/j.psyneuen.2019.02.007

33. Striepens N, Kendrick KM, Maier W, et al. Prosocial effects of oxytocin and clinical evidence for its therapeutic potential. Front Neuroendocrinol 2011; 32:426–450. https://doi.org/10.1016/j.yfrne.2011.07.001

34. Guzmán YF, Tronson NC, Jovasevic V, et al. Fear-enhancing effects of septal oxytocin receptors. NatNeurosci 2013: 16:1185–1187. https://doi.org10.1038/nn.3465

35. Shamay-Tsoory SG, Fischer M, Dvash J, et al. Intranasal administration of increases envy and schadenfreude (gloating). Biol Psychiatry 2009; 66:864–870. https://doi.org/10.1016/j.biopsych.2009.06.009

36. Goodson JL, Thompson RR. Nonapeptide mechanisms of social cognition, behaviour and species-specific social systems. Curr Opin Neurobiol 2010; 20:784–794. https://doi.org/10.1016/j.conb.2010.08.020.

37. Bredewold R, Smith CJ, Dumais KM, et al. Sex-specific modulation of juvenile social play behavior by vasopressin and depends on social context. Front Behav Neurosci 2014; https://doi.org/10.3389/fnbeh.2014.00216

